# Pharmacophore-Inspired Virtual Receptor: Generating Realistic Peptide Binding Data for Machine Learning

**DOI:** 10.64898/2026.11.16.766906

**Authors:** Kilian Conde-Frieboes, Carsten Stahlhut

## Abstract

VReceptor is a virtual receptor simulation framework that generates realistic, sequence-dependent binding data for peptide-based therapeutics. It combines pharmacophore-inspired weighting schemes with amino acid similarity metrics to evaluate peptide design strategies in a controlled environment. Validation using the Prolactin-Releasing Peptide (PrRP) dataset confirms VReceptor’s ability to approximate binding behaviours and optimize experimental planning. This framework empowers researchers in computational peptide design and active learning workflows while high-lighting areas for further refinement.

## 1 Introduction

Peptide-based therapeutics have demonstrated significant potential over the past few decades for the treatment of various diseases with both specificity and biological safety [1]. The development of *in silico* tools for drug discovery has revolutionised early-stage pharmaceutical research, enabling rapid hypothesis testing and virtual screening of vast molecular libraries. However, a persistent challenge remains: how to validate these computational models effectively, especially when experimental resources are limited or prohibitively expensive. This is particularly true for peptide-based therapeutics, where the combinatorial explosion of possible sequences makes exhaustive experimental testing infeasible.

To address this, we introduce VReceptor, a virtual receptor simulation framework designed to generate realistic, sequence-dependent binding affinity data. Inspired by the pharmacophore concept originally proposed by Paul Ehrlich [2, 3], VReceptor provides a controlled and interpretable environment for evaluating peptide design strategies and active learning algorithms. It is particularly suited for benchmarking machine learning models in peptide optimisation tasks, where the goal is to predict or improve binding affinity based on sequence information.

Unlike purely random or black-box data generators, VReceptor is built on a transparent and biologically motivated model. It simulates ligand–receptor interactions by assigning importance to specific amino acid positions using a pharmacophore-like weighting scheme. These weights are then combined with amino acid distance metrics to produce a binding score that reflects both sequence composition and structural relevance. The result is a synthetic yet biologically plausible dataset that can be used to test hypotheses, compare design strategies, and train predictive models.

### 1.1 Related work

To address the challenge of molecular optimization, recent research has introduced the Poli library, which provides a suite of artificial fitness functions, or oracles, designed to serve as reliable proxies for molecular traits of interest [4, 5]. While this library facilitates systematic benchmarking of molecular optimization methods specifically for protein optimization, further efforts are needed to broaden the range of modalities to include peptides and to validate the consistency and reliability of these oracles across various relevant wet-lab characterizations.

Although numerous bioinformatics tools such as PeptideRanker [6] exist for targeting the modulation of bioactive peptides, [7] highlights a general deficiency in their effectiveness. A significant challenge is the weak correlation between predictions made by these tools and experimentally derived IC50 values. This discrepancy indicates that many models may not adequately capture the complex biological interactions governing peptide activity, making them less reliable for practical applications. This underscores the urgent need for more accurate computational methods to enhance the predictive validity of bioinformatics tools, facilitating their integration into peptide design and leading to more effective bioactive peptides. Similarly, recent benchmarking work, as exemplified in [8], provides benchmarking framework tailored for protein folding neural networks in protein-peptide complex prediction. While AlphaFold 3 [9] accordingly to [8] demonstrates strong performance in structure prediction, they note that the confidence metrics correlate poorly with experimental binding affinities.

In this context, VReceptor offers a complementary approach: a transparent and controllable virtual environment for generating realistic synthetic data to accelerate method development. Although we validate the realism of VReceptor by fitting it to experimentally measured affinities from the PrRP peptide family [10], this serves only as a demonstration of plausibility rather than the primary contribution of the work. This work makes the following contributions:

- **A realistic virtual receptor framework (VReceptor):** A transparent, pharmacophore-inspired simulation model that generates biologically plausible, sequence-dependent binding affinity data for peptides.
- **A synthetic, controllable environment for benchmarking and accelerating peptide-model development:** VReceptor enables systematic testing of peptide optimisation strategies, search algorithms, active learning methods, and machine learning models under realistic and interpretable conditions—supporting rapid iteration before experimental investment.
- **A biologically interpretable receptor model:** The framework combines position-specific weighting with amino acid distance metrics, offering mechanistic transparency absent in black-box oracle generators.
- **Empirical demonstration of realism:** We show that VReceptor can be calibrated to approximate real binding landscapes using data from the PrRP peptide family—serving as a validation step, not the main contribution.

## 2 Peptide-Receptor Binding Model

The foundation of the VReceptor framework is a generative model that simulates peptide–receptor binding affinities based on peptide sequence composition. The core idea is to construct a function that maps a peptide sequence to a predicted binding strength, mimicking the behaviour of a biological receptor. This function is designed to be deterministic and sequence-dependent, ensuring that similar sequences yield similar binding profiles, thereby enabling meaningful comparisons and learning.

The binding affinity is modelled as a function of the peptide sequence (eq.1):

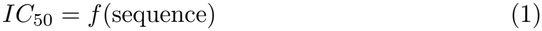

To achieve this, the model begins with a reference peptide of length *n*, where each position *i* ∈ {1, …, *n}* is assigned an importance score *b*_*i*_. These scores reflect the contribution of each position to the overall binding interaction and are derived from a hypothetical pharmacophore model. This model is expressed as a mixture of Gaussian functions *g*_*k*_ (eq. 2, Fig. 1).

**Figure 1:**
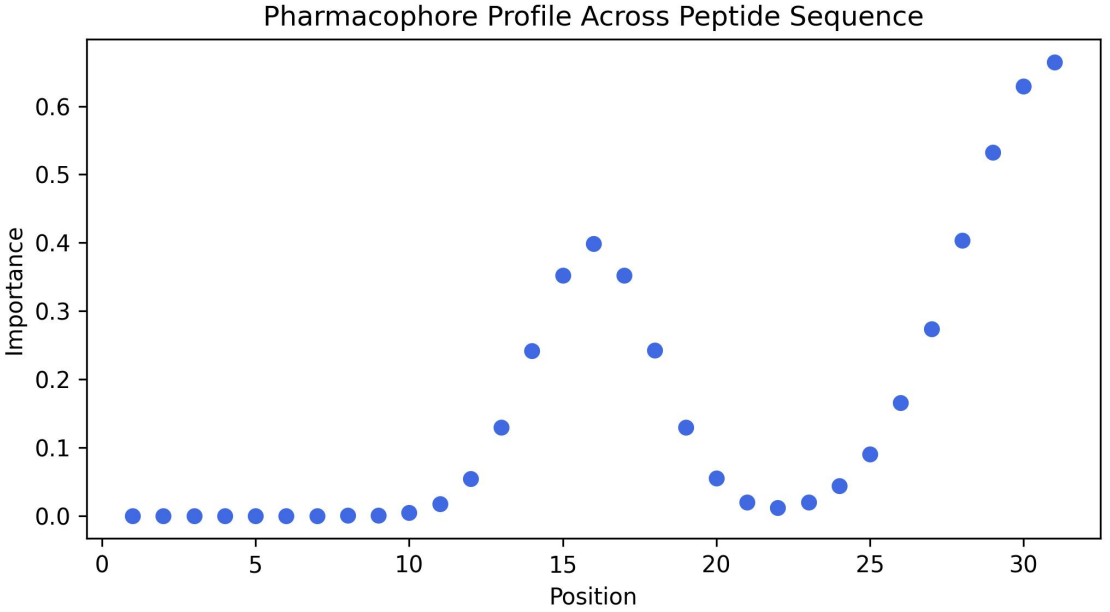
Hypothetical pharmacophore model for a peptide with 31 amino acids.

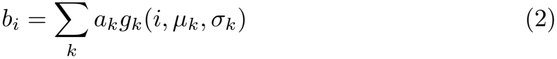

Here, each Gaussian *g*_*k*_ is parameterised by an amplitude *a*_*k*_, a mean *µ*_*k*_, and a width *σ*_*k*_, allowing the model to flexibly encode position-specific relevance across the peptide. The selection of gaussian functions to model the importance, or weights, of different amino acid stretches in the peptides allows a smaller number of parameter as compared to individual weights, but also reflects ligand-receptor interactions, where small areas in the peptide interact with the receptor, but nearby amino acids have a diminishing effect.

In the VReceptor model, each amino acid is represented by a feature vector 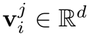, where *d* denotes the number of physicochemical descriptors used—herein derived from z-scales as described in [11], *i* indicates the position in the peptide and *j* indicates the amino acid selected from the amino acid alphabet. These descriptors capture properties such as hydrophobicity, steric bulk, and electronic characteristics.

To calculate the score of a target peptide relative to the reference peptide, the model needs a measure of distance between the amino acids. The distance function 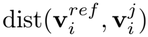 quantifies the dissimilarity between the amino acid at position *i* in the reference peptide and the amino acid in the same position of the target peptide based on their physicochemical properties (e.g., z-scale descriptors), and is elaborated in Section 2.1.

Next, a weight matrix **W** ∈ ℝ^*n×m*^ is constructed, where *m* is the number of amino acids in the alphabet (typically 20). Each element *W*_*ij*_ represents the weighted contribution of substituting the amino acid at position *i* with amino acid *j*, and is computed as (eq. 3):

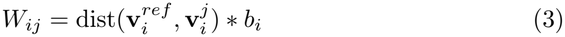

In the VReceptor model, each peptide sequence is represented using a one-hot encoding matrix **E**, which captures the identity of amino acids at each position in the sequence. For a peptide of length *n* and an amino acid alphabet of size *m* (typically the 20 standard amino acids), the matrix **E** ∈ {0, 1}^*n×m*^ is defined such that:

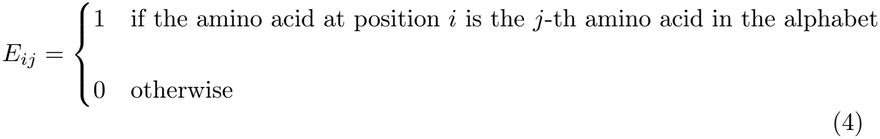

Each row of **E** corresponds to a position in the peptide, and each column corresponds to a specific amino acid. Only one entry per row is set to 1, indicating the presence of a particular amino acid at that position, while all other entries are 0.

This encoding allows the model to compute the *Hadamard product* **R** ∈ ℝ^*n×m*^ (eq. 5), where **W** is the weight matrix derived from the pharmacophore model and distance scores. The resulting matrix **R** reflects the contribution of each amino acid to the overall binding affinity of the target peptide, which is then summed to produce a scalar prediction score.

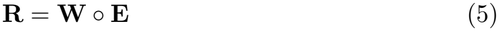

To simulate experimental variability, homoscedastic noise is added to the computed binding affinity. The final expression for the relative binding affinity is (eq. 6):

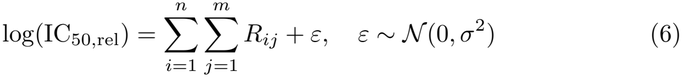

Here, *R*_*ij*_ denotes the weighted contribution of amino acid *j* at position *i*, and *ε* is a normally distributed noise term with zero mean and variance *σ*^2^, consistent with the homoscedasticity assumption commonly used in linear mixed models for assay variability estimation [12].

Together, these components form the basis of the VReceptor simulation engine, enabling the generation of realistic, interpretable binding data for downstream analysis and model training. The model does not account for interactions between amino acids within the peptide nor any structural or conformational considerations; it focuses solely on position-specific importance scores.

### 2.1 Distance Measures

Three different distance measures have been integrated into the VReceptor, with the possibility of further extensions. Similarly, extension of the VReceptor can be done with respect to the feature vector **v** by replacing z-scale derived physicochemical descriptors with alternative representative descriptors.

#### 2.1.1 Cosine Distance

To assess the distance between amino acids based on their vector representations (e.g. z5 descriptors), we define a pairwise distance measure using the cosine of the angle between their feature vectors. Let us denote the vector of descriptors of amino acids *g* as **v**_*g*_ and the corresponding vector of amino acid *h* as **v**_*h*_. The cosine distance is then defined as (eq. 7):

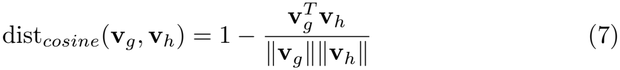

This formulation yields values in the interval [0, 2], where higher values indicate greater distance. It captures the angular proximity between amino acid descriptors and is particularly useful when the direction of the feature vector is more informative than its magnitude.

#### 2.1.2 Chebyshev-like Distance

To quantify distance based on the maximum absolute difference between features, we define a Chebyshev-like distance. Let **v**_*g*_ be the descriptor vector of amino acid *g* and **v**_*h*_ be the feature vector of amino acid *h*. The raw Chebyshev distance is the following with *d* being the dimension of the z-scale vectors **v** (eq. 8):

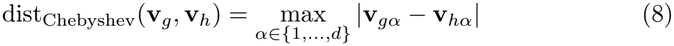

To normalize this distance across the amino acid set, we divide by the maximum observed difference for each feature (eq. 9).

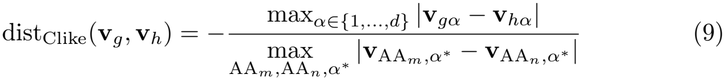

This normalized form ensures comparability across features and amino acid pairs, making it suitable for models where dominant feature differences are most relevant.

#### 2.1.3 L2 Distance

To quantify the distance between amino acids based on their physicochemical properties, we define a pairwise distance measure derived from the L2 norm of normalized feature vectors. Let **v**_*g*_ denote the vector of properties of amino acid *g*, in this case the z-scale vector. Each vector is normalized to the unit length (eq. 10):

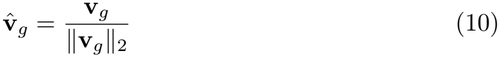

The distance between amino acids *g* and *h* is then defined as (eq. 11):

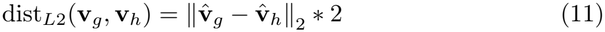

This formulation ensures that the distance values are in the interval [0, 2], with higher values indicating greater distance. The approach captures relative proximity in the normalized feature space and is particularly suited for comparing amino acids in embedded or kernel-based models. Figure 2 shows a comparison of the three different distance measures.

**Figure 2:**
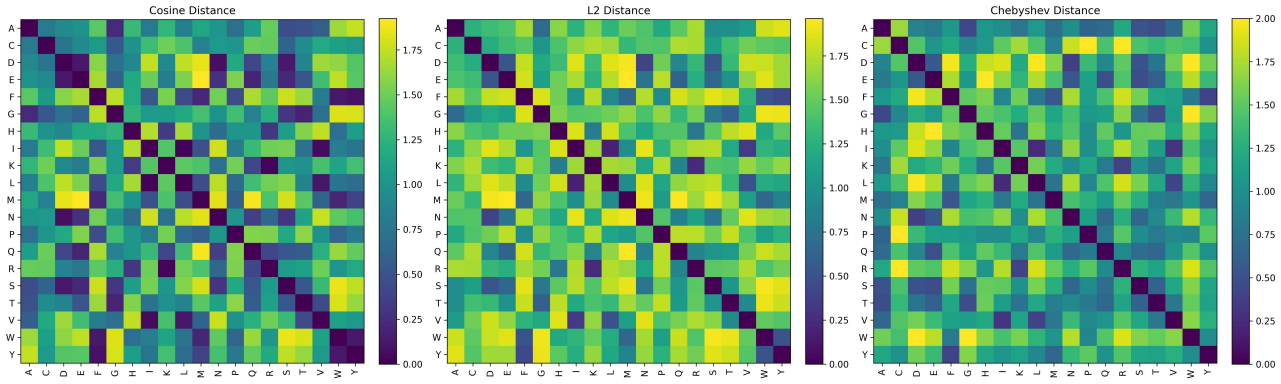
Visual comparison between the 3 discussed distance measures.

## 3 Experiments

The intended purpose of this tool was to serve as a data generator for evaluating peptide library design strategies, tasks that would otherwise be prohibitively expensive and time-consuming to evaluate experimentally. A simple question is, for example, whether a full mutational scan of a given target peptide should be conducted or whether a random library of peptides with multiple mutations would yield a better model. All experiments were conducted on compute architecture featuring an x86_64 configuration with a single Intel Xeon Platinum 8360H CPU @ 3.00 GHz. In the supplied notebook (see GitHub) we address this problem for a peptide of length 31. We chose the Prolactin-Releasing Peptide (PrRP, [10]) sequence as a starting peptide and created an artificial pharmacophore with the use of two Gaussian functions as shown in figure 1. A complete mutational scan with the 20 canonical amino acids yields 589 peptides (31* 19). We compared this design with a random design of equal size. Sequences from the 30 million possible triple mutations for this target peptide were selected. After generating simulated data with the primed VReceptor, two similar models are trained on these datasets and evaluated on a random test set of 200 peptides, again selected from the set of all triple mutated peptides (figure 3).

**Figure 3:**
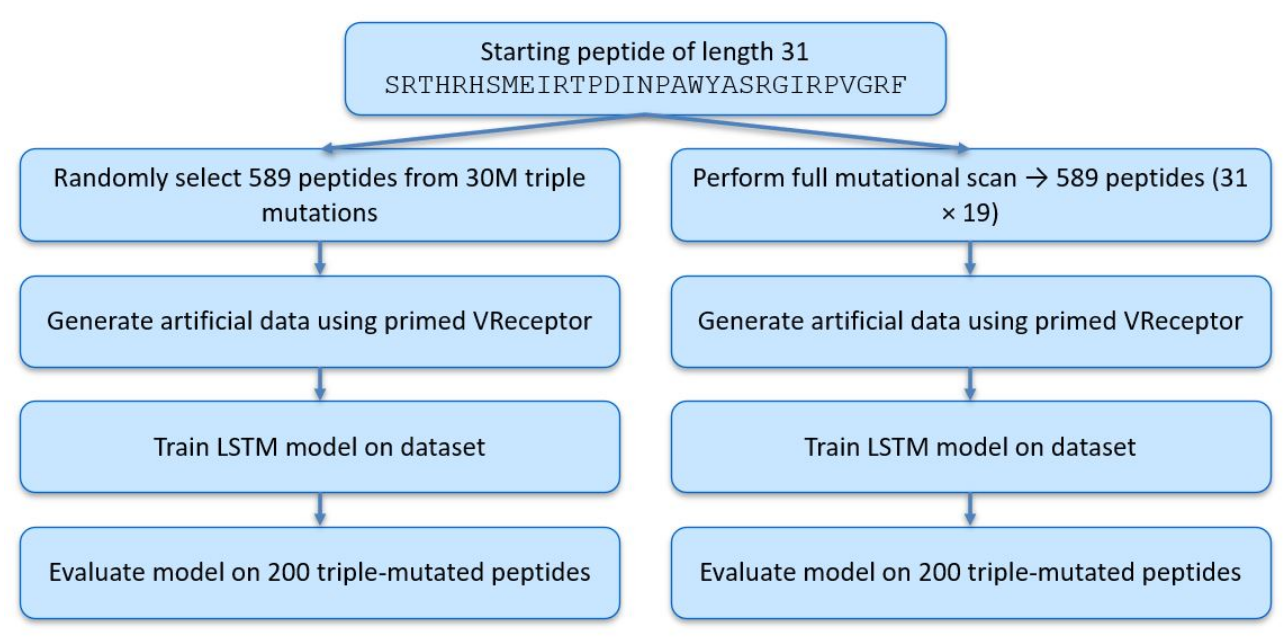
Flow chart to compare two different design strategies using the VReceptor.

The models consist of a 2-layer, bidirectional LSTM [13] neural network as the regression model. The hidden state of the LSTM layer is fed into the output layer. After training the models are evaluated on the test data set. In this case the model trained on the random data set has a smaller mean-squared error as compared to the systematic generated scan data set (0.023 vs 0.140) (fig. 4).

**Figure 4:**
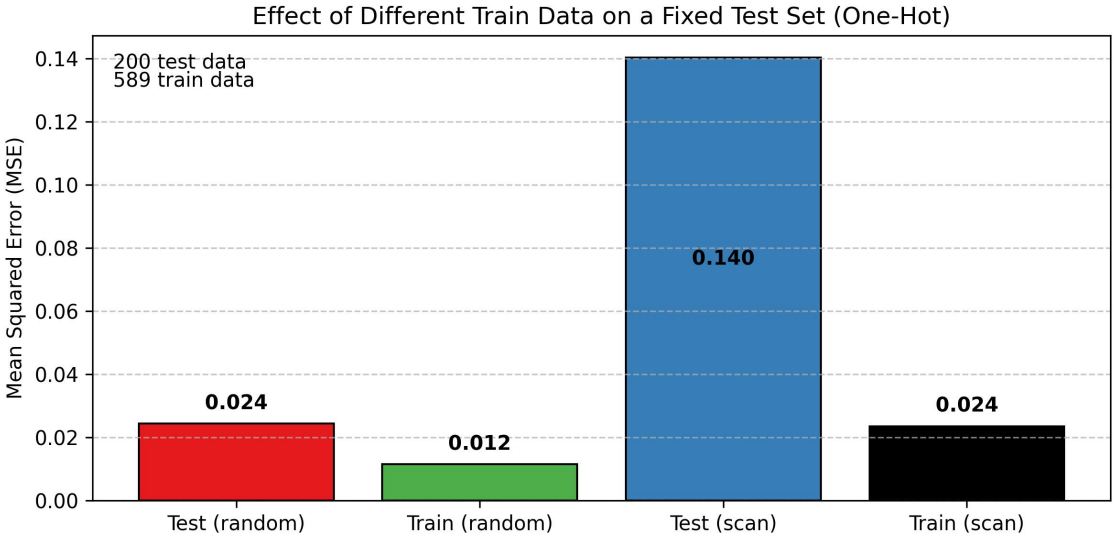
Train/Test error for the scan and random design strategy

Figure 4 shows the mean squared error (mse) comparing the two different models.

### 3.1 Realism of VReceptor

During the development and testing of VReceptor, a central question emerged: How realistic is this model? Can peptide–receptor interactions truly be captured with such simplicity? VReceptor is intentionally designed with a limited number of parameters, determined by the number of Gaussian functions *N*_*G*_ specified during initialization. The only additional degree of freedom lies in the choice of distance measure dist(*g, h*), which governs how amino acid substitutions are evaluated. To assess the model’s authenticity and practical value, we explored fitting these minimal parameters to real experimental data. This approach allows us to evaluate whether VReceptor can meaningfully replicate observed binding behaviours, thereby validating its usefulness as a simulation tool.

#### 3.1.1 The PrRP dataset

Prolactin-Releasing Peptide (PrRP) is a neuropeptide involved in the regulation of food intake and energy homeostasis [10]. It exerts its physiological effects by binding to at least three G-protein coupled receptors: GPR10, NPFF receptor 1 (NPFFR1), and NPFF receptor 2 (NPFFR2). The dataset used to validate the VReceptor tool comprises experimentally measured binding affinities (log *IC*_50_ values) of PrRP-derived peptides against these three receptors. The listed data are relative to wildtype human PrRP. All peptides in the dataset are 31 amino acids long and C-terminally amidated, reflecting their biologically active form. A subset of peptides includes fatty acid modifications, denoted by an ‘X’ in the sequence; however, these were excluded from the present analysis to maintain consistency in molecular structure and avoid confounding effects. The estimated standard deviation is 0.1 on the log_10_ scale.

#### 3.1.2 Parameter Fitting Procedure for VReceptor Models

To evaluate the performance of the VReceptor model under different amino acid similarity metrics, we implemented a parameter optimisation workflow using three distinct similarity functions: cosine similarity, Chebyshev-like distance, and L2 norm-based similarity. For each similarity function, we trained a VReceptor model to predict the binding affinity (log *IC*_50_) of peptide sequences using a minimal set of parameters. The optimisation objective was to minimise the mean squared error (MSE) between predicted and experimentally measured log *IC*_50_ values, represented as *ŷ*_*n*_ and *y*_*n*_, respectively, as detailed in (eq. 12):

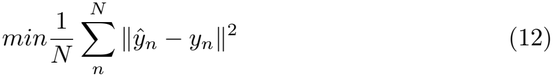

We selected a subset of the PrRP dataset where the target response variable (relGPR10) was below a threshold of 2.0. The data was split into training and test sets using a 50:50 ratio with a fixed random seed for reproducibility. The model parameters were optimised using the scipy.optimize.minimize function. The parameter vector included the positions and amplitudes of two Gaussian functions used to define the pharmacophore model. A grid of initial parameter guesses was explored to avoid local minima, iterating over combinations of Gaussian centres and amplitudes.

For each candidate parameter set, a VReceptor instance was constructed and used to simulate binding affinities for the training sequences. The predicted values were compared to experimental data using MSE as the loss function. The best-performing parameter set was retained for each similarity function. The optimised models were then evaluated on both training and test sets. Predicted log *IC*_50_ values were compared to experimental values using Pearson and Spearman correlation coefficients, and mean squared error (MSE). The results were stored for each similarity function and visualised to compare model performance across metrics.

When comparing the VReceptor model metrics with a Ridge linear regression with the same data split and an empirical determined alpha = 10, the VReceptor performs similar good (figure 5, table 1). The larger difference between the VReceptor and Ridge regression on the train data is most likely a reflection of the vast difference in parameter, 6 vs 155, respectively, and a train set size of 250 compounds.

**Table 1:** Performance metrics for different methods on training and test sets.

| **Method** | **Pearson r** | **Spearman r** | **MSE** |
|---|---|---|---|
| **Training Set** |  |  |  |
| Cosine | 0.82 | 0.80 | 0.131 |
| Chebyshev | 0.85 | 0.83 | 0.109 |
| L2_sim | 0.85 | 0.83 | 0.108 |
| Ridge | **0.95** | **0.92** | **0.036** |
| **Test Set** |  |  |  |
| Cosine | 0.77 | 0.75 | 0.155 |
| Chebyshev | **0.83** | **0.80** | **0.118** |
| L2_sim | **0.83** | **0.80** | **0.118** |
| Ridge | 0.80 | 0.73 | 0.140 |

**Figure 5:**
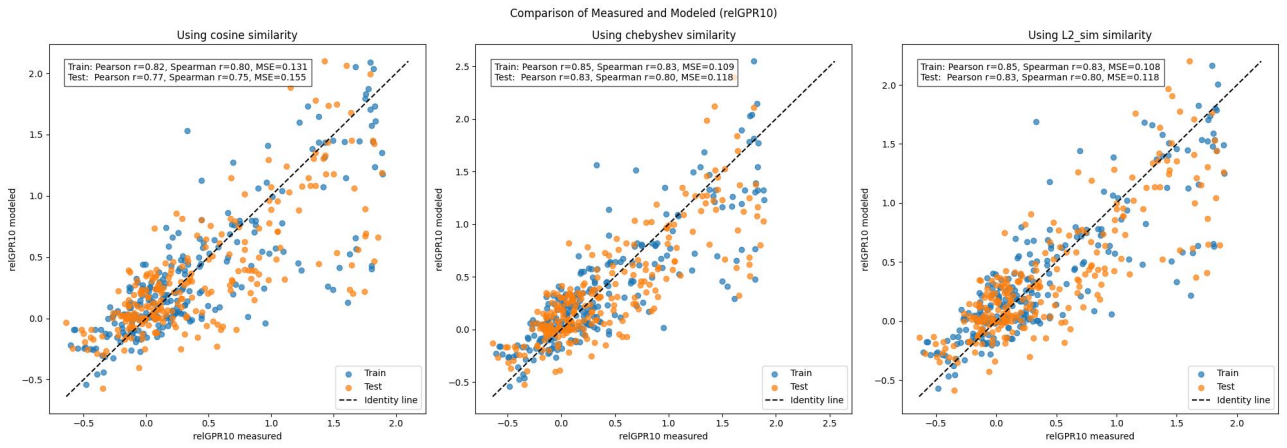
Performance of VReceptor models using three different similarity measures

The fitted VRecptors can predict the test data with decent metrics. The model using the cosine distance is slightly worse than the other two, but still acceptable.

## 4 Discussion

The VReceptor framework presents a streamlined yet biologically grounded approach to simulating peptide–receptor interactions. By combining a pharmacophore-inspired weighting scheme with interpretable amino acid similarity metrics, it enables the generation of synthetic binding data that reflects meaningful sequence dependencies. This makes it particularly valuable for benchmarking peptide design strategies and evaluating active learning workflows.

A key strength of VReceptor is its balance between simplicity and flexibility. The model is parameterised by a small number of Gaussian functions, which define the positional importance of amino acids in a peptide sequence. This allows for intuitive control over the simulated binding landscape while maintaining computational efficiency. The integration of multiple distance measures—cosine, Chebyshev-like, and L2—further enhances the model’s adaptability, enabling users to explore how different assumptions about amino acid similarity influence predictive performance.

The application to the PrRP dataset demonstrates that VReceptor can approximate experimental binding affinities with reasonable accuracy, even when using a minimal parameter set. This suggests that the model can serve as a practical surrogate for experimental assays in early-stage research, where rapid iteration and hypothesis testing are critical. The comparison of design strategies—full mutational scans versus random triple-mutant libraries—also highlights the tool’s utility in guiding experimental planning.

However, the model’s simplicity also imposes limitations. It assumes additive contributions from individual amino acid positions and does not account for higher-order interactions or structural constraints. While this is a deliberate design choice to preserve interpretability, it may limit the model’s applicability in more complex biological contexts. This can be seen when fitting a model on the other two provided receptor data (NPFFR1 and NPFFR2). These fittings have higher mean-squared error values and lower Spearman and Pearson correlations. Two possible explanations are the collected data are not diverse enough to provide a good data set and/or there are other effects like e.g. the influence of the helicity of the peptides on binding affinities which are not captured in this simple model. Also the fitting of the model to real data was done to evaluate the realism of VReceptor and the simple global minimum search strategy might not be thorough enough.

## 5 Conclusion & Future Directions

In summary, VReceptor offers a lightweight, interpretable, and extensible platform for simulating realistic peptide binding data. It is well-suited for use in computational peptide design, active learning evaluation, and methodological benchmarking. Its realism and applicability is validated through approximating the binding behavior of a real wetlab experimental dataset with enhanced generalization compared to simple ridge regression when facing limited experimental data.

Beyond its role as a synthetic data generator and benchmarking tool, VReceptor can serve as a robust baseline in predictive machine learning workflows. Its transparent, biologically motivated design allows researchers to systematically evaluate the performance of more complex models against a well-understood reference. By incorporating additional features—such as structural priors or higher-order sequence interactions—VReceptor may achieve predictive accuracy on par with state-of-the-art machine learning approaches, while retaining interpretability and computational efficiency.

We gratefully acknowledge Claus Bekker Jeppesen and Lone Honoré at Novo Nordisk for their invaluable contribution in collecting the PrRP dataset. KCF and CS were employees of Novo Nordisk A/S at the time the work was conducted. KCF remains an employee of Novo Nordisk A/S.

